## Supplementary figures and images for "Clearance of protein aggregates during cell division"

### Supplemental figures

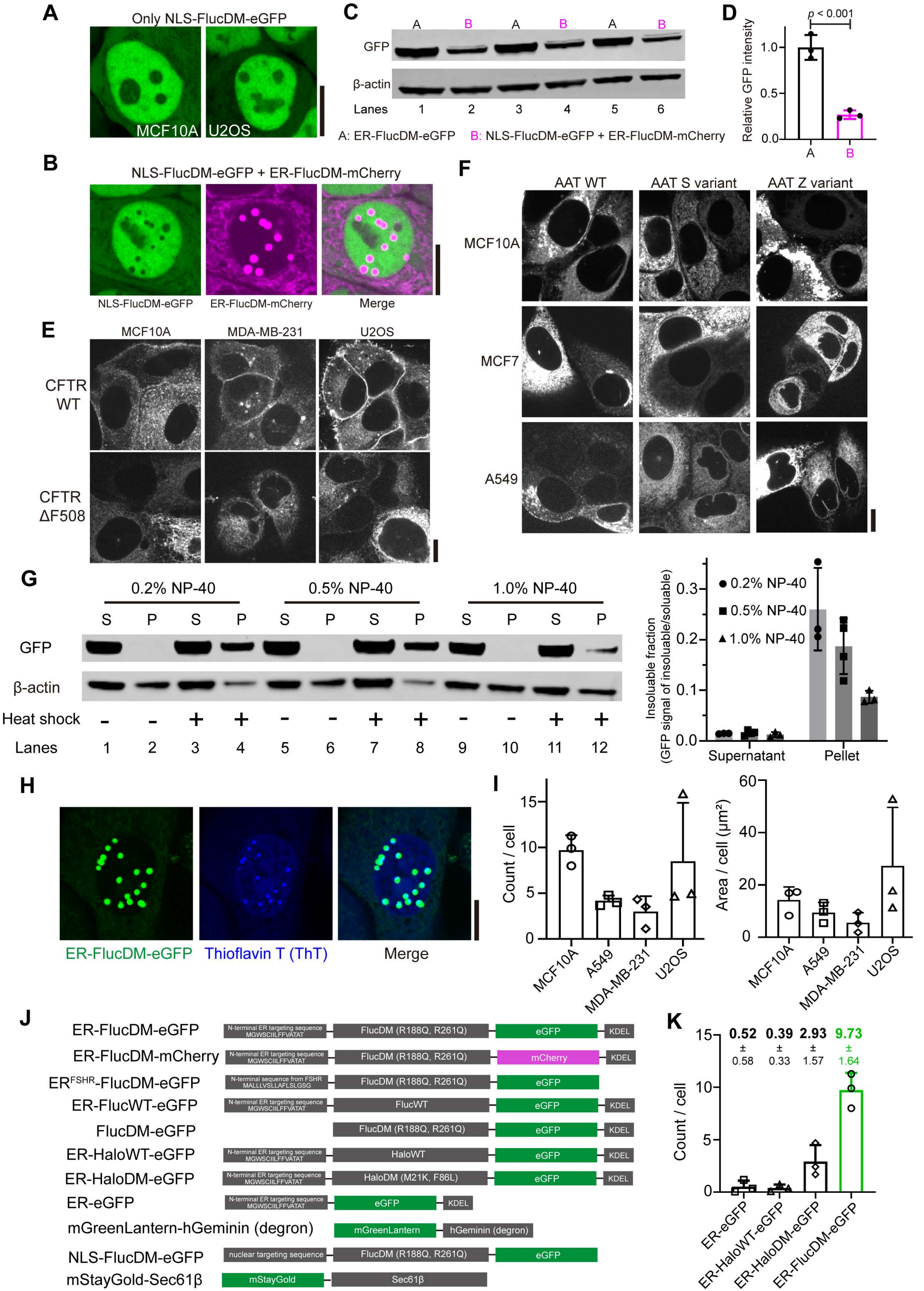

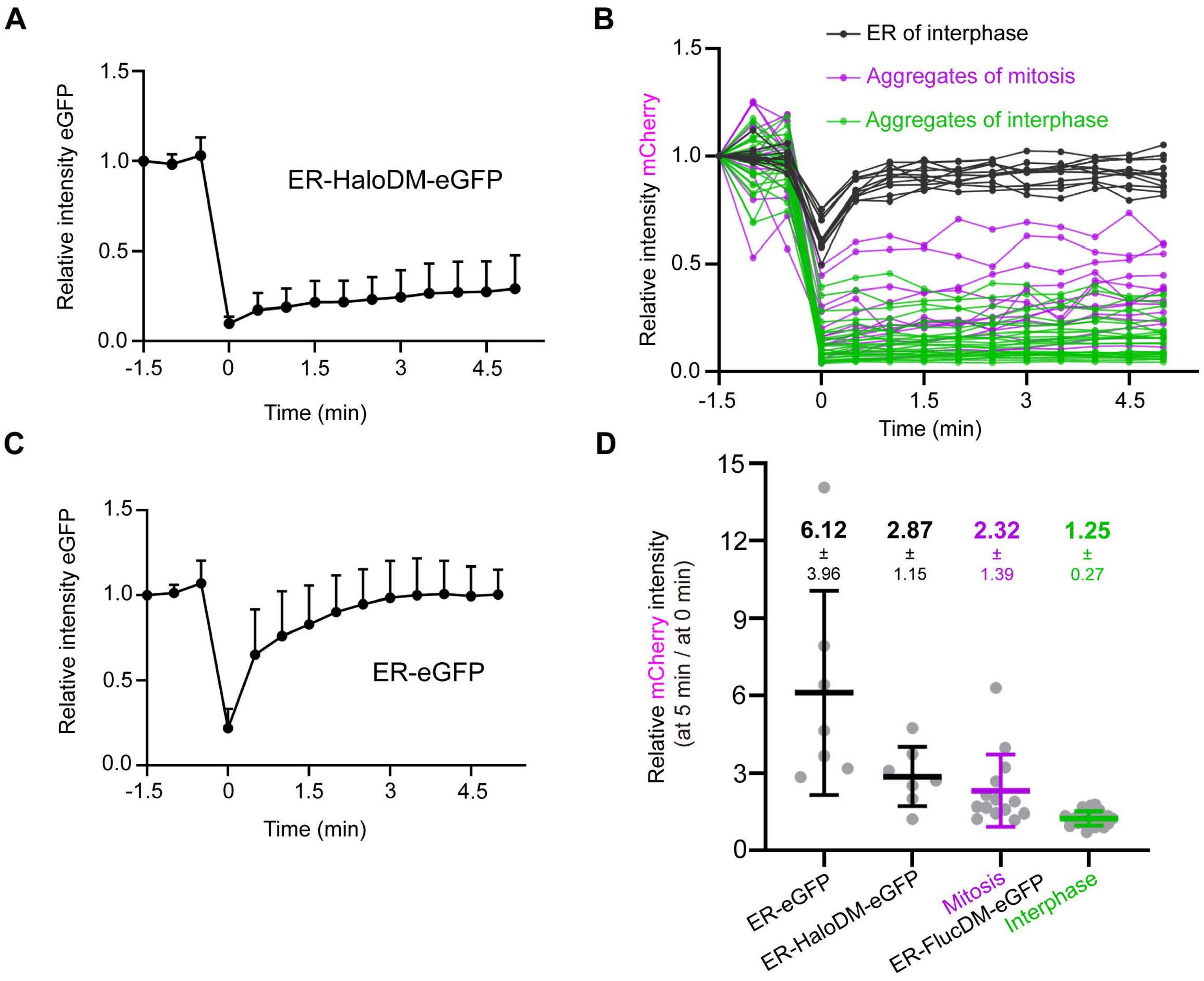

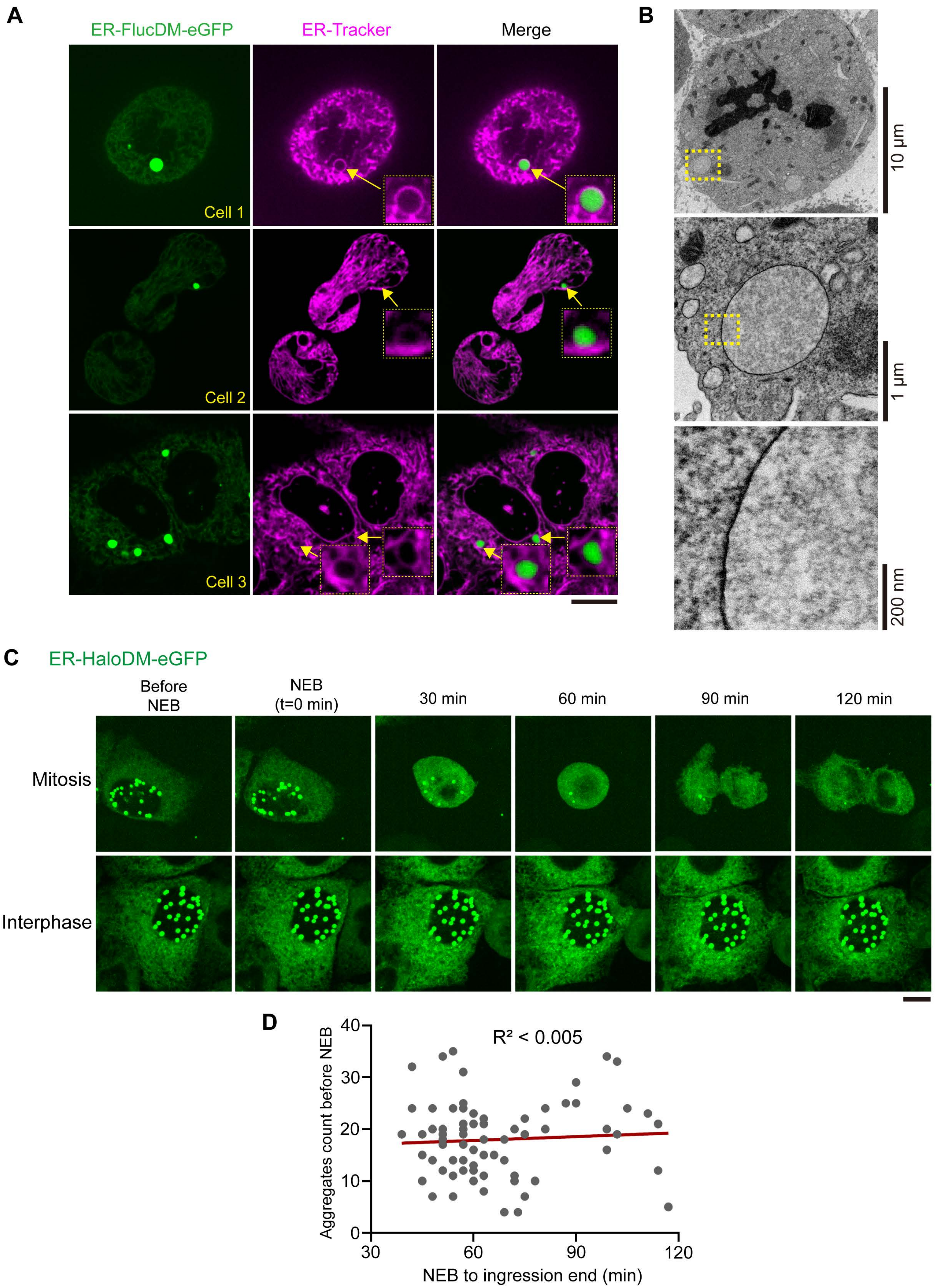

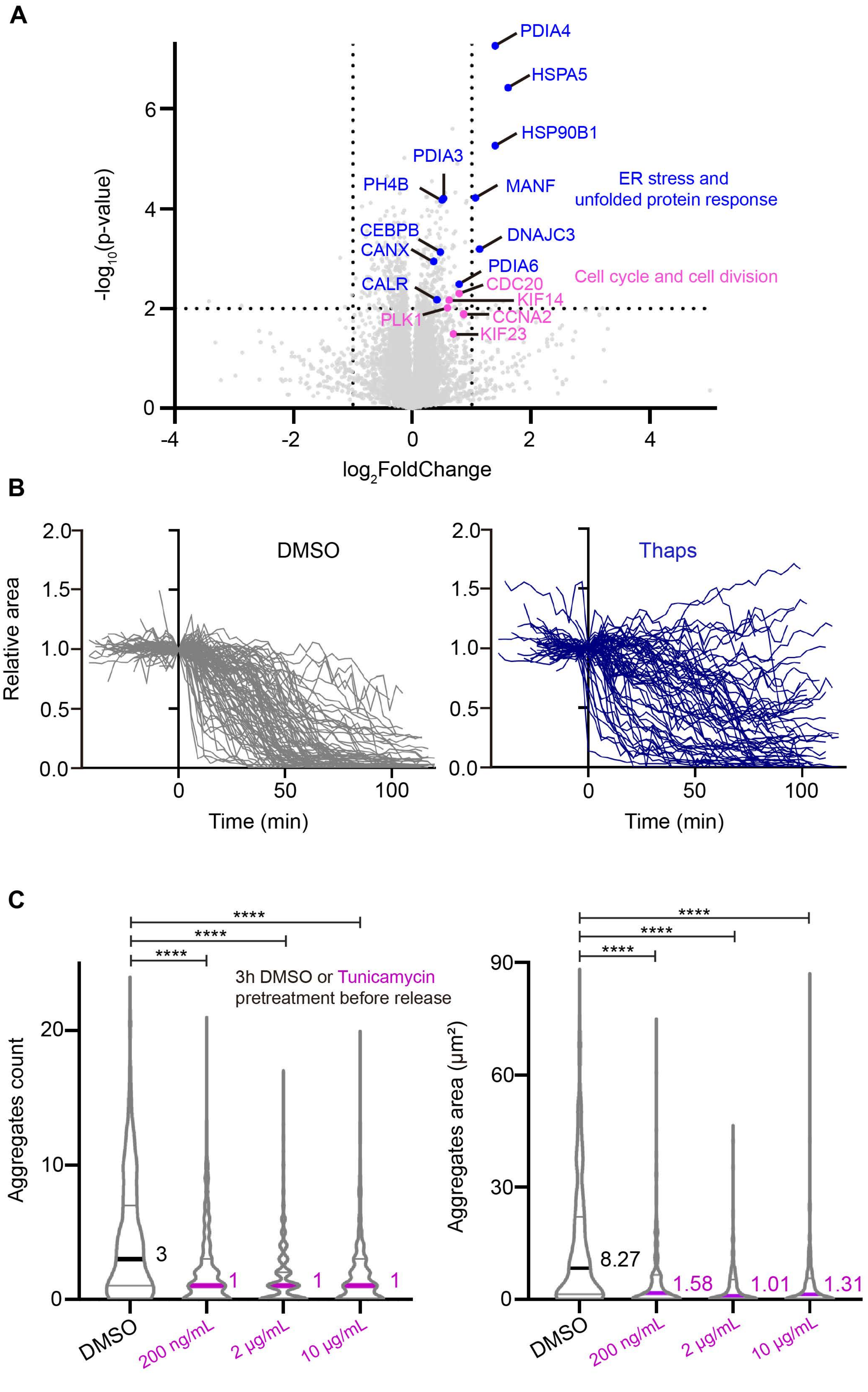

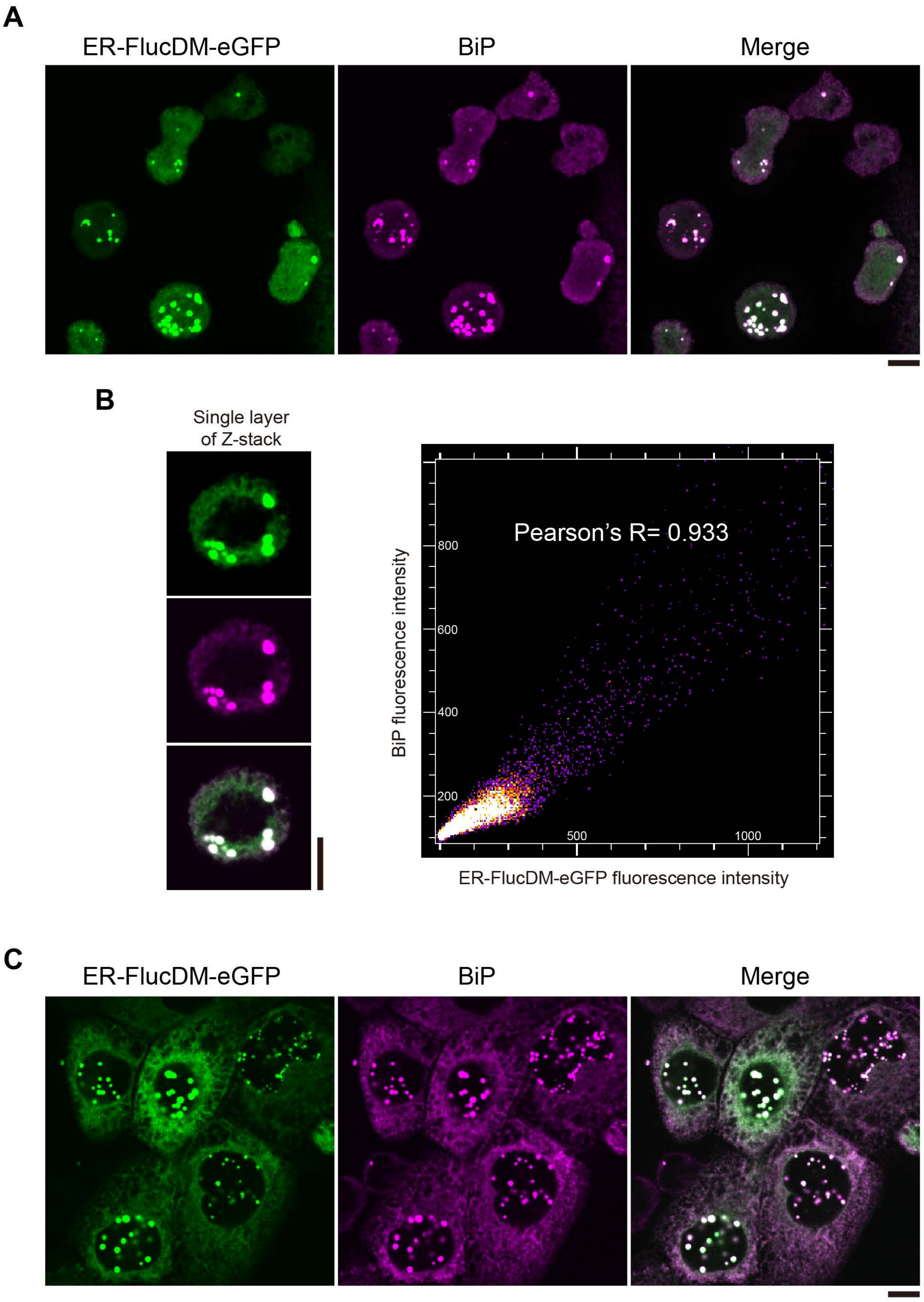

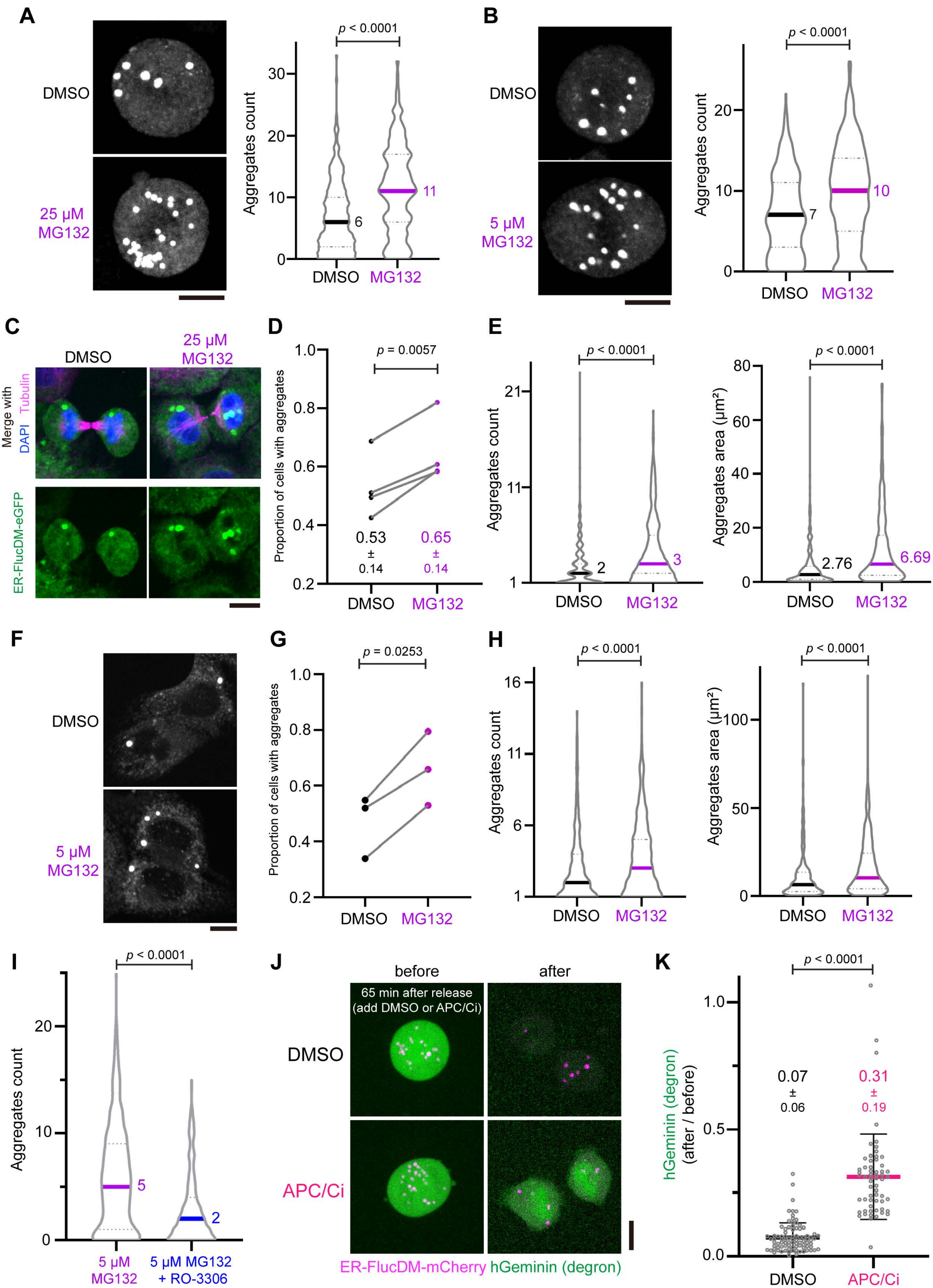

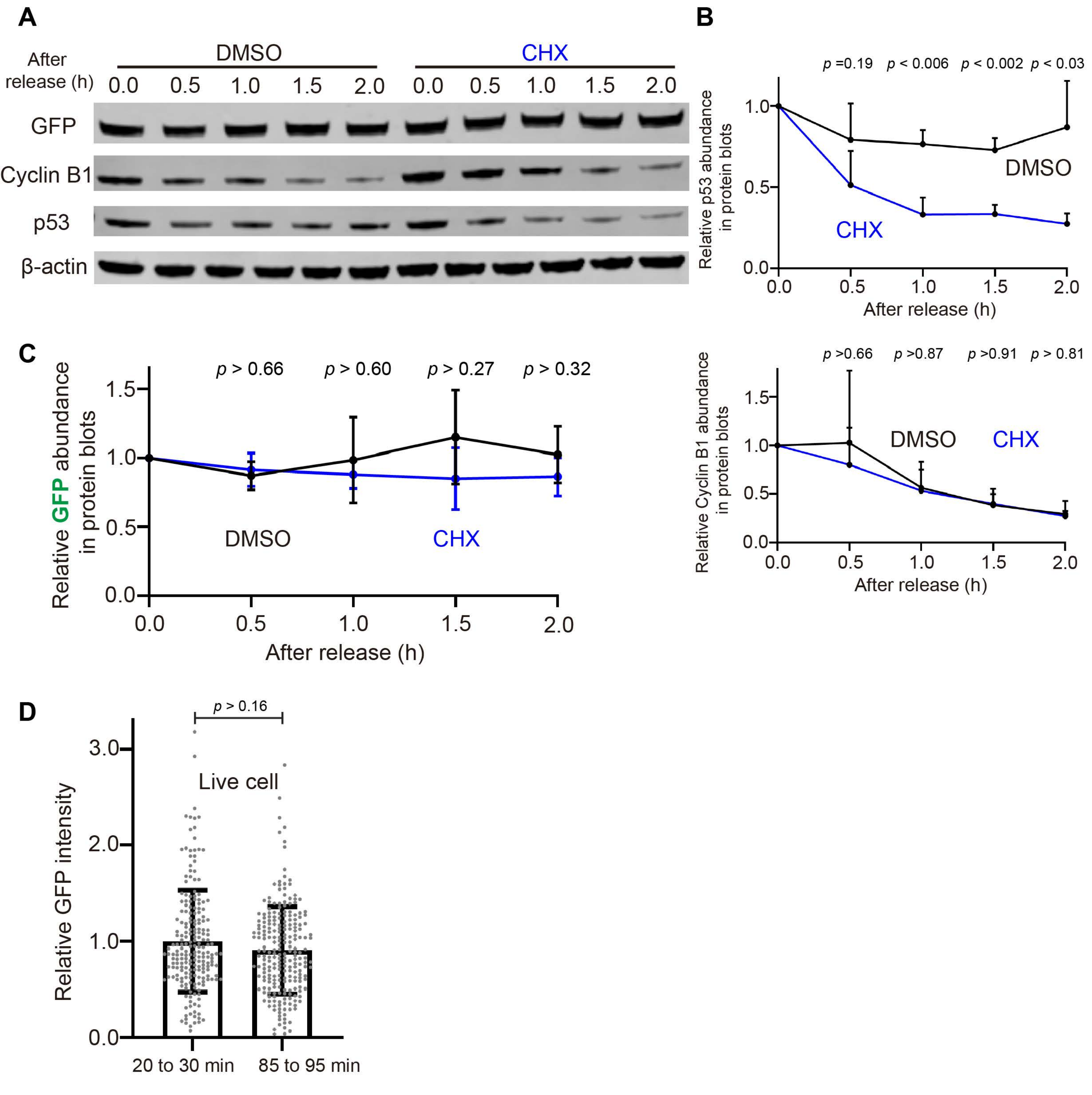
